# Bidirectional hybridization between *Ulva prolifera* and *U. linza* (Ulvophyceae, Chlorophyta): Evidence for compatibility and paternal chloroplast inheritance

**DOI:** 10.64898/2026.07.18.739365

**Authors:** Zhengzhao Xu, Jin Zhao, Peng Jiang

## Abstract

*Ulva prolifera* and *U. linza* are closely related species, with abundant adult thalli and reproductive cells co-occurring extensively in time and space during the Yellow Sea green tides. Elucidating their hybridization compatibility is crucial for species delimitation, assessing interspecific gene flow, and evaluating the ecological impacts of green tides. Previous studies suggested asymmetric gamete compatibility (only *U. prolifera* mt^+^ × *U. linza* mt^−^), but lacked sex-linked markers to reliably identify hybrid diploids and their reproductive modes, and did not examine chloroplast inheritance. Here, we performed bidirectional crosses using sexual strains of different geographic origins from both parents, with sex-linked markers to quantify progeny genotypes, examine fertility and reproduction pathway of F_1_ hybrids, and trace chloroplast inheritance using the species-specific *pet*B marker. Our results showed that: (1) F_1_ hybrids were obtained in both cross directions, with significantly higher frequency in the direct cross (*U. linza* mt^+^ × *U. prolifera* mt^−^), indicating no complete reproductive isolation in either direction; the biased compatibility likely reflected genetic background differences among the limited strain combinations in a single study. (2) A considerable number of germinated progeny arose from parthenogenesis of parental gametes. (3) F_1_ hybrids from both crosses could undergo meiosis to form gametes and develop into gametophytes; additionally, F_1_ from the reciprocal cross produced diploid spores for asexual reproduction, suggesting meiotic disturbance. (4) Chloroplasts were maternally inherited in selfing of *U. prolifera* parent, but in all F_1_ hybrids they were paternally inherited, indicating a potential reversal of the inheritance pattern due to interspecific hybridization. These findings provided new insights into the potential for genetic exchange between *U. prolifera* and *U. linza*.

## 1. Introduction

Species of *Ulva* are widely distributed in coastal, estuarine, and brackish-water environments worldwide and constitute an important group of primary producers in coastal ecosystems (Shimada et al., 2008). They generally exhibit efficient nutrient uptake, rapid growth, and high reproductive potential. Under favorite environmental conditions, especially eutrophication, some species can proliferate rapidly and accumulate as extensive floating or suspended biomass, thereby forming green tides with substantial adverse effects on coastal ecosystems (Duong et al., 2025). To date, more than 100 species are currently recognized within the genus according to AlgaeBase (https://www.algaebase.org). Due to the facts that *Ulva* thalli are structurally simple with limited number of diagnostic morphological characters which are unstable to environments (Hofmann et al., 2010), species delimitation and phylogenetic reconstruction have increasingly relied on molecular markers. Such efforts have not only revealed cryptic species that are morphologically similar but also identified several genetically closely related lineages (Hofmann et al., 2010; Steinhagen et al., 2019). To address the challenge that existing molecular markers are insufficient to discriminate among these closely related taxa (Shimada et al., 2008; Kang et al., 2019), it has been proposed that species boundaries within *Ulva* should be evaluated not only through morphological and molecular approaches, but also through artificial crossing experiments that directly assess reproductive compatibility among closely related taxa (Hiraoka et al., 2017; Tran et al., 2022).

*U. prolifera* and *U. linza* are representative closely related taxa and provide an important system for investigating species-boundary formation in *Ulva*. Multiple phylogenetic studies have shown that almost all molecular markers applicable to *Ulva*, including ITS, *rbc*L, and *tuf*A, have grouped them into a single phylogenetic clade (Wang et al., 2010; Saunders & Kucera, 2010; Leliaert et al., 2009). Only the highly variable 5S rDNA non-transcribed spacers (5S-NTS) can separate them into two relatively independent genetic groups (Shimada et al., 2008). Despite the high genetic similarity suggesting that the two may have undergone relatively recent speciation, they have already exhibited certain differentiation in ecological distribution and morphological traits: *U. prolifera* commonly occurs in estuarine and brackish-water habitats and has relatively conspicuous branching, whereas *U. linza* is mainly distributed in marine intertidal zones and is usually sparsely branched or unbranched (Cui et al., 2018). Therefore, compared with genetic divergence, the more pronounced morphological differentiation was considered likely to be related to their long-term adaptation to contrasting environments. However, during large-scale green tides in the Yellow Sea, abundant adult thalli and reproductive cells of the two species co-occur extensively in both space and time (Zhang et al., 2019; Han et al., 2020), theoretically creating substantial opportunities for natural interspecific hybridization. Previous studies have assessed their compatibility using artificial interspecific crosses between sexual strains of different geographic origins, and found that gametes of female *U. prolifera* can fuse with those of male *U. linza*, whereas no gamete fusion was observed in the reciprocal cross (Hiraoka et al., 2011). Xie et al. (2020) further reported that the F_1_ hybrid sporophytes were fertile and their offspring could develop into adults.

However, the absence of mating-type-specific molecular markers at the time made it difficult to definitively confirm, quantify, or track certain aspects of the hybridization process. First, the verification of F_1_ hybrids remained challenging, relying primarily on microscopic observation of gamete fusion (Hiraoka et al., 2011; Cui et al., 2018; Xie et al., 2020). *Ulva* exhibit a typical isomorphic alternation of generations, wherein haploid gametophytes produce gametes of different mating types; upon fusion, these form diploid sporophytes, which subsequently undergo meiosis to produce meiospores that germinate into male and female gametophytes. It should be noted that following hybridization, germinated progeny may represent either F_1_ hybrid sporophytes (2n) arising from gamete fusion between the two parental sexes (with unknown germination rates), or male and female gametophytes (n) resulting from parthenogenetic development of gametes; however, sporophytes and gametophytes are morphologically indistinguishable. Consequently, the proportion of F_1_ hybrid sporophytes among germinated seedlings—a critical indicator reflecting hybridization compatibility—cannot be accurately determined. Second, the fertility assessment of F_1_ hybrid sporophytes remains insufficient. Studies in *U. prolifera* and *U. linza* have demonstrated that their sporophytes (2n) can either undergo meiosis to form meiospores that germinate into gametophytes (n), or produce diploid asexual spores that germinate into sporophytes (2n) identical to themselves (Innes, 1987; Hiraoka et al., 2003; Ichihara et al., 2019). Which reproductive pathways are adopted by F_1_ hybrid sporophytes? Furthermore, do differences exist between direct and reciprocal cross combinations? These questions bear significant implications for understanding whether hybridization events may exert long-term evolutionary impacts; however, they currently remain entirely unknown.

In addition to nuclear genome recombination, the selective inheritance of organellar genomes also constitutes a critical determinant of the genetic composition in F_1_ hybrids. The chloroplast genome encodes genes essential for photosynthesis and other key metabolic processes, and its parental origin not only directly determines the organellar genetic background of F_1_ hybrids, but may also modulate gene expression and metabolic coordination through cytonuclear interactions, thereby potentially affecting offspring growth performance, environmental adaptability, and ecological fitness (Sloan et al., 2018). Therefore, elucidating the parental origin of chloroplasts in F_1_ hybrids is essential for a comprehensive understanding of the genetic consequences of interspecific hybridization. In most angiosperms and algal taxa, including *Ulva*, chloroplasts are generally inherited maternally (Reboud & Zeyl, 1994; Kagami et al., 2008; Miyamura, 2010; Choi et al., 2020; Liu, 2022), although paternal, biparental, and stochastic inheritance involving the retention of chloroplasts from either parent have also been reported (Mogi et al., 2009; Zhang & Sodmergen, 2010). Studies in angiosperms have further shown that chloroplast inheritance patterns in interspecific hybrids may differ from those observed within the parental species (Kumari et al., 2011; Li et al., 2013; Shrestha et al., 2021). However, the parental origin and inheritance stability of chloroplasts during interspecific hybridization in *Ulva*, as well as the consistency of these patterns between bidirectional crosses, remain poorly understood. Elucidating chloroplast inheritance in F_1_ hybrids is therefore important for understanding the genetic consequences of interspecific hybridization in *Ulva* and its potential ecological implications.

In this study, *U. prolifera* and *U. linza* were used as parental species, with gametophytic strains representing different geographic origins and mating types selected for bidirectional artificial crossing experiments. Sex-linked molecular markers were utilized to systematically evaluate the cross compatibility between these two closely related species, detect the genetic origin of germinated progeny, and examine the fertility of F_1_ hybrid sporophytes and their capacity for meiotic completion. In addition, chloroplast inheritance patterns were tracked and identified across generations, spanning from the parental strains through the F_1_ hybrid sporophytes to their descendants. This study will provide novel insights into the genetic exchange potential between *U. prolifera* and *U. linza*.

## 2. Materials and methods

### 2.1 Algal materials and culture conditions

*Ulva prolifera* O.F.Müller and *U. linza* Linnaeus strains were used as the P generation in hybridization experiments. These were unialgal gametophytic cultures maintained in our laboratory. Information on sample collection and genetic identification for all algae strains were presented in Table 1. Genomic DNA was extracted from each individual using a Plant Genomic DNA Extraction Kit (Tiangen Biotech Co., Ltd., Beijing, China) following the manufacturer’s instructions. Two molecular markers, ITS and 5S-NTS (Leskinen and Pamilo, 1997; Shimada et al., 2008), were combined to use for species-level identification. All primers used in this study were shown in Table 2.

**Table 1.** Collection information and mating type of gametophytic strains used as the P generation.

| Species | Collection date | Location | Sample ID | Mating type |
| --- | --- | --- | --- | --- |
| *U. prolifera* | 29/07/2007 | Qingdao, Shandong, China | S096-1<sup>a</sup> | mt<sup>+</sup> |
| *U. prolifera* | 29/07/2007 | Qingdao, Shandong, China | S096-2<sup>a</sup> | mt<sup>-</sup> |
| *U. prolifera* | 27/04/2024 | Xiamen, Fujian, China | U687-1 | mt<sup>-</sup> |
| *U. prolifera* | 27/04/2024 | Xiamen, Fujian, China | U687-2 | mt<sup>+</sup> |
| *U. prolifera* | 27/04/2024 | Xiamen, Fujian, China | U687-3 | mt<sup>+</sup> |
| *U. linza* | 13/12/2024 | Yantai, Shandong, China | 24YT12-1 | mt<sup>-</sup> |
| *U. linza* | 22/05/2023 | Qingdao, Shandong, China | 23QD07-1<sup>b</sup> | mt<sup>+</sup> |
| *U. linza* | 22/05/2023 | Qingdao, Shandong, China | 23QD07-2<sup>b</sup> | mt<sup>+</sup> |
| *U. linza* | 22/05/2023 | Qingdao, Shandong, China | 23QD07-3<sup>b</sup> | mt<sup>-</sup> |

<sup>a</sup> Prepared from the meiospore of sporophytic strain S096.
<sup>b</sup> Prepared from the meiospore of sporophytic strain 23QD07.

**Table 2.**
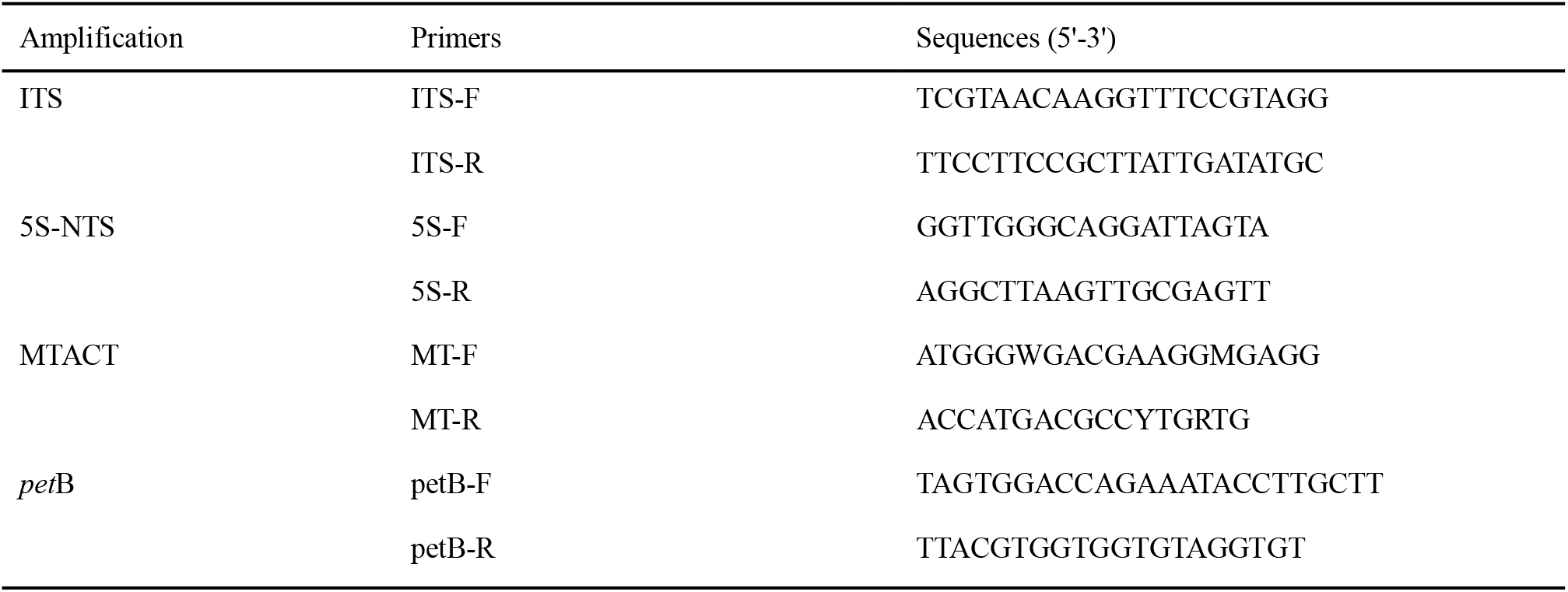
All primers used in this research.

The mating type of the samples was detected using the MTACT molecular marker targeting the mating type locus in *Ulva* (Xu, 2024). PCR amplification was conducted using a T960 thermal cycler (Heal Force, Shanghai, China). The amplified products were detected by 1.5% agarose gel electrophoresis, and the Trans2K Plus II DNA Marker (TransGen Biotech Co., Ltd., Beijing, China) was employed in this study. Haploid female gametophytes (mt^+^) produced an amplification fragment of approximately 340 bp, while male gametophytes (mt^−^) yielded one of approximately 210 bp; owing to heterozygosity at this locus in diploid individuals, both sizes of amplification fragments appeared simultaneously in the sporophyte. Consequently, the generation or sexual characteristics of the samples could be readily distinguished after gel electrophoresis of the PCR products.

In order to maintain or accelerate the proliferation of algal strains, individual algal samples were cultured in the light incubator (model GXZ-380B, Ningbo Southeast Instrument Co., Ltd., Ningbo, China) using von Stosch’s enriched (VSE) medium, with weekly medium replacements. Thalli were maintained under vegetative growth conditions at 16°C, a photoperiod of 12 h light : 12 h dark, and photosynthetic irradiance of approximately 100 µmol photons m^-2^ • s^-1^.

### 2.2 Interspecific hybridization

In this study, both direct and reciprocal crosses were conducted between the two species. The direct cross comprised female *U. linza* × male *U. prolifera*, and the reciprocal cross comprised female *U. prolifera* × male *U. linza*. The preparation of male and female gametes was based on the established protocols with appropriate modification (Hiraoka et al., 2003; Liu et al., 2022a). Gametophyte parents of both sexes were sectioned into 1–2 mm-long thallus segments using a sterile scalpel, transferred to new Petri dishes with VSE medium, and incubated under conditions of 20°C, photoperiod of 14 h light : 10 h dark, and a photosynthetic photon flux density of approximately 150 µmol photons m^-2^ • s^-1^ to induce gametogenesis and gamete release. Three replicates were set up for each of the hybrid combinations. Following gamete discharge, the Petri dishes were moved to a gradient light field generated by a unilateral light source. Taking advantage of the positive phototaxis exhibited by the gametes, the accumulated gametes along the illuminated side were gently aspirated using a micropipette. 50 µL each of female and male gamete suspensions were rapidly mixed; the mixture was then incubated statically for 12 h to facilitate gamete fusion and zygote formation, after which the Petri dishes were transferred to the normal culture conditions described above for continued cultivation.

For the germinated seedlings from each interspecific hybridization group, once they had grown to approximately 2–4 cm, a subset of individual plants were selected. Genomic DNA was extracted from each alga individually as described previously, and PCR amplification of the MTACT molecular marker was performed to identify the F_1_ hybrid diploids.

### 2.3 Assessment of meiotic competence in F_1_ hybrid diploids

Individual algal seedlings identified as F_1_ hybrid diploids from both the direct and reciprocal cross groups were selected for continuous cultivation. After the seedlings have developed into adult thalli, gametogenesis and gamete release were induced using the aforementioned method to assess the fertility of the F_1_ hybrid diploids. Subsequently, the germinated F_1_-derived progeny were likewise identified using the MTACT molecular marker to evaluate the meiotic capability of the F_1_ hybrid diploids.

### 2.4 Morphological observation

Observations of thallus reproductive structures and released reproductive cells were conducted under a BH2 light microscope (Olympus Corp., Tokyo, Japan). Lugol’s iodine was used to stain the swimming zoids (Xie et al., 2020). All images of morphological features were taken with a CCD camera (Scope Tek MDC200, Mingshi, Ningbo, China) mounted on the microscope and the ScopePhoto 3.0 software.

### 2.5 Tracking chloroplast inheritance from P generation through F_1_ hybrid diploids to F_1_-derived progeny

In order to determine the genetic origin of chloroplast transmission from parents to hybrid progeny, a fragment of chloroplast *pet*B gene was amplified across P generation, F_1_ hybrid diploids and F_1_-derived progeny. The PCR amplification procedure for *pet*B consisted of an initial denaturation at 94°C for 10 min, followed by 35 cycles of 94°C for 1 min, 54°C for 50 s, and 72°C for 1 min, with a final extension at 72°C for 10 min. The amplified products exhibited a reproducible length difference between the two parents. A ∼150 bp product was generated in *U. linza*, whereas a ∼1.8 kb fragment containing one additional intron was produced in *U. prolifera* (Liu et al., 2022b).

## 3. Results

### 3.1 Reconfirmation of the parental mating type

Before initiating the hybridization experiment, the mating type of each parental pair was reconfirmed by PCR using the MT-F/R primers of the MTACT molecular marker for all nine parental gametophyte strains of the two species. The agarose gel electrophoresis results of the amplified products were shown in Fig. 1, which were fully consistent with the identification results listed in Table 1.

**Fig. 1.**
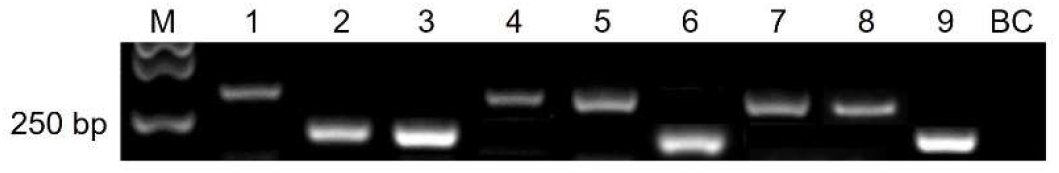
Genotyping of all parental strains by PCR amplification of the MTACT marker. M: DNA marker; Lane 1–5: *U. prolifera* parents S096-1, S096-2, U687-1, U687-2, and U687-3, respectively; Lane 6–9: *U. linza* parents 24YT12-1, 23QD07-1, 23QD07-2, and 23QD07-3, respectively; BC: blank control.

### 3.2 Hybridization between U. prolifera and U. linza

Interspecific hybridization was performed according to the design shown in Table 3. A total of 10 cross groups were established, comprising four direct crosses and six reciprocal crosses, using nine female and male gametophyte parents from *U. prolifera* and *U. linza*.

**Table 3.**
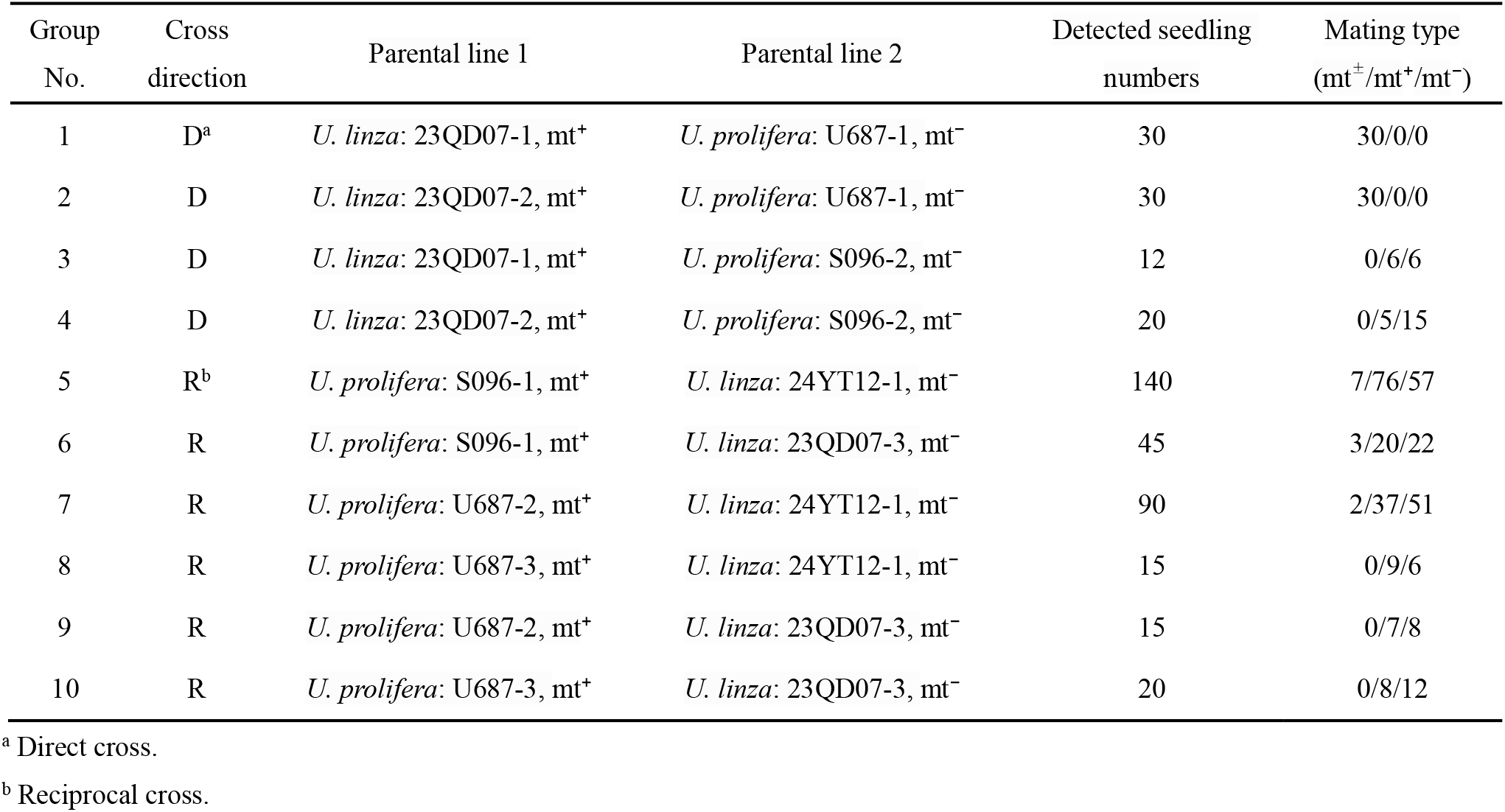
Hybrid combinations and detected mating types in germinated progeny.

Gametophyte parents at the vegetative growth stage were selected (Fig. 2a). After approximately 48–72 h of induction by thallus sectioning, microscopic examination revealed that all parental individuals had successively differentiated to form gametangia (Fig. 2b). Upon unilateral light irradiation, the released reproductive cells in all groups were observed to aggregate at the edge of the Petri dish on the light-facing side, indicating positive phototaxis (Fig. 2c). Results of Lugol’s staining showed that all were biflagellate gametes (Fig. 2d). Equal volumes of freshly released female and male gametes were mixed, and gamete fusion was observed under the microscope within minutes (Fig. 2e). After approximately 14 d of continued culture, germinated progeny approximately 0.5 cm in length were visible in all hybridization combinations (Fig. 2f).

**Fig. 2.**
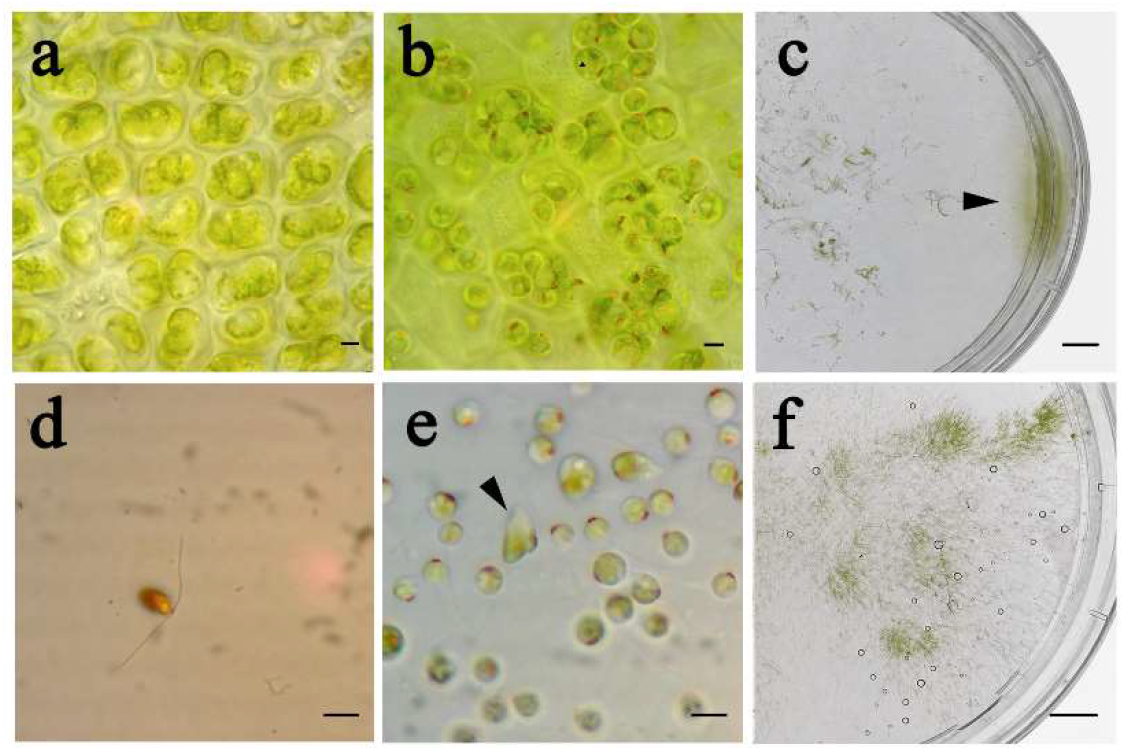
Interspecific hybridization process between *U. prolifera* and *U. linza*. (a) Gametophyte parent of P generation before thallus sectioning. (b) Differentiation and formation of the gametangia. (c) The black arrowhead points to reproductive cells aggregated due to positive phototaxis. (d) Biflagellate gamete. (e) The black arrowhead points to fusing gametes. (f) Germinated progeny. Scale bar = 5 µm (a, b, d, e) or 1 cm (c, f).

Once the seedlings have grown to 2–4 cm, a subset of samples was selected from each cross group and genotyped for the MTACT molecular marker using the MT-F/R primers. The electrophoresis results for amplified products were shown in Fig. 3. Based on the identified mating types (Table 3), it was shown that the seedlings included F_1_ hybrid diploids (mt^±^) as well as haploid gametophytes (mt^+^/mt^−^) derived from parthenogenesis of female and male gametes, respectively. Among the four direct cross groups, F_1_ hybrid diploids were detected only in groups 1 and 2 (Fig. 3a), with a detection rate of 100% (n = 30) in both; no F_1_ hybrid diploids were detected in groups 3 and 4, and all seedlings originated from parthenogenesis of both female and male gametes. Among the six reciprocal cross groups, F_1_ hybrid diploids were detected only in groups 5–7 (Fig. 3b), but at markedly lower proportions than in the direct cross groups, ranging from merely 2.2% to 6.6%; no F_1_ hybrid diploids were detected in groups 8–10, and all seedlings originated from parthenogenesis of both female and male gametes. These results indicated that F_1_ hybrid sporophytes could be detected in both cross directions, although a stronger gamete compatibility bias toward the direct cross was observed in our study.

**Fig. 3.**
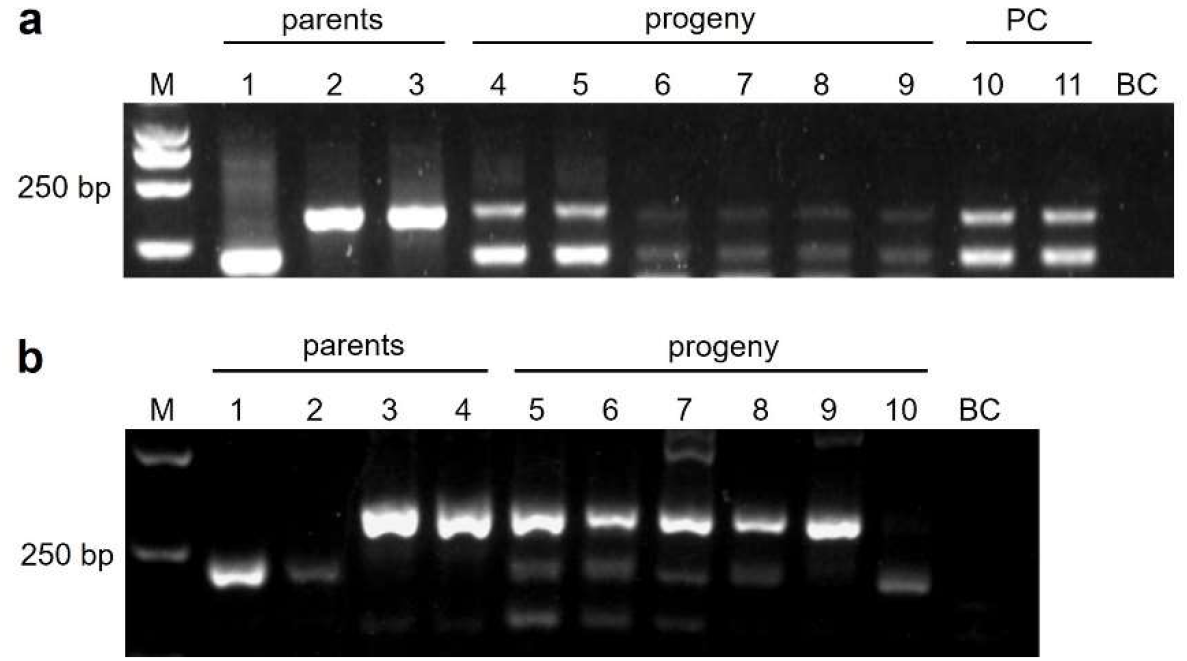
PCR-detection for MTACT marker in parents and germinated progeny. (a) Direct cross groups 1 and 2. M: DNA marker; Lane 1: *U. prolifera* parent U687-1; Lane 2–3: *U. linza* parents 23QD07-1 and 23QD07-2; Lane 4–6: F_1_ hybrid diploids from cross group 1; Lane 7–9: F_1_ hybrid diploids from cross group 2; Lane 10–11: progeny of S096-1×S096-2; PC: positive control; BC: blank control. (b) Reciprocal cross groups 5, 6 and 7. M: DNA marker; Lane 1–2: *U. linza* parents 24YT12-1 and 23QD07-3; Lane 3–4: *U. prolifera* parents S096-1 and U687-2; Lane 5–8: F_1_ hybrid diploids from cross groups 5, 5, 6, and 7, respectively; Lane 9: parthenogenetic progeny of *U. prolifera* mt^+^ parent S096-1; Lane 10: parthenogenetic progeny of *U. linza* mt^−^ parent 24YT12-1; BC: blank control.

### 3.3 Assessment of meiotic competence in F_1_ hybrid diploids

To investigate whether the F_1_ hybrid diploids were fertile and capable of meiosis, one F_1_ hybrid diploid individual (mt^±^, 2n) was selected from each of the direct cross groups 1 and 2 and from the reciprocal cross group 5, respectively. Sporangium formation was induced through thallus sectioning, and the progeny generated by released reproductive cells were subsequently subjected to MTACT-PCR assay. The results showed that after 2–3 days of induction, thallus fragments in all groups turned yellow and, upon microscopic examination, were found capable of forming sporangia and releasing reproductive cells.

The MTACT-PCR assay results for the germinated F_1_-derived seedlings were shown in Fig. 4. For the F_1_-derived progeny from direct cross groups 1 and 2, the mt^±^:mt^+^:mt^−^ ratio was 0:2:3 (n = 5) in each group; all seedlings were haploid gametophytes (Fig. 4a), with a female-to-male ratio approximating 1 : 1. For the F_1_-derived progeny from reciprocal cross group 5, the mt^±^:mt^+^:mt^−^ ratio was 6:0:19 (n = 25); six diploid sporophytes were detected, while all remaining seedlings were haploid male gametophytes, with no haploid female gametophytes identified (Fig. 4b). These results indicated that while meiosis appeared to proceed normally in F_1_ diploids from the direct cross, it was severely impaired in those from the reciprocal cross.

**Fig. 4.**
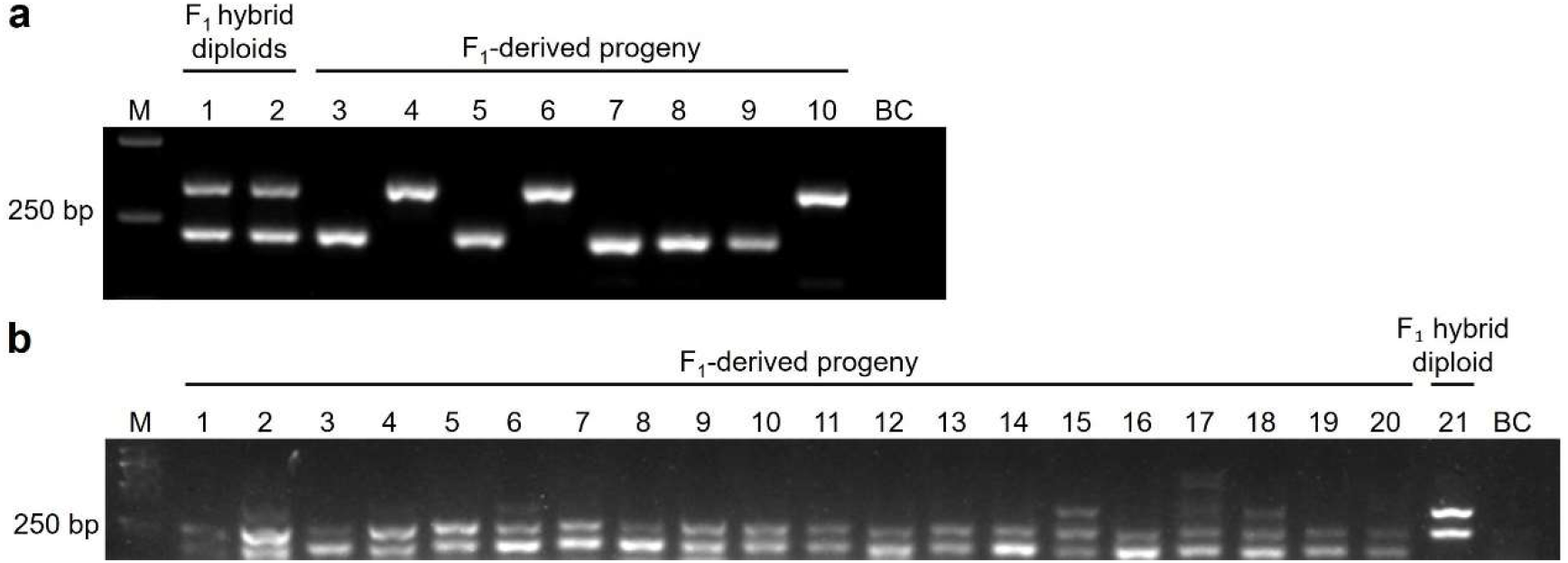
PCR-detection for MTACT marker in F_1_ hybrid diploids and their progeny. (a) Direct cross groups 1 and 2. M: DNA marker; Lane 1–2: F_1_ hybrid diploids from cross groups 1 and 2; Lane 3–6: F_1_-derived haploids (mt^+^/mt^−^) from cross group 1; Lane 7–10: F_1_-derived haploids (mt^+^/mt^−^) from cross group 2; BC: blank control. (b) Lane 1–20: F_1_-derived male haploids (mt^−^) and apomictic diploids (mt^±^) from cross group 5; Lane 21: F_1_ hybrid diploid from cross group 5; BC: blank control.

### 3.5 Chloroplast inheritance pattern

To track the transmission pattern of chloroplasts from the P-generation parents to the F_1_ hybrid sporophytes and their progeny, the chloroplast *pet*B marker was used for tracing and detection.

First, in the P generation, six parents from direct cross groups 1 and 2 and reciprocal cross groups 5 and 6 that were capable of producing F_1_ hybrid sporophytes were subjected to *pet*B detection, including strains U687-1 (mt^−^) and S096-f (mt^+^) from *U. prolifera*, as well as strains 23QD07-1 (mt^+^), 23QD07-2 (mt^+^), 24YT12-1 (mt^−^), and 23QD07-3 (mt^−^) from *U. linza*. The amplification results matched expectations, yielding ∼1.8 kb fragments in *U. prolifera* and ∼150 bp fragments in *U. linza* (Fig. 5a, b).

**Fig. 5.**
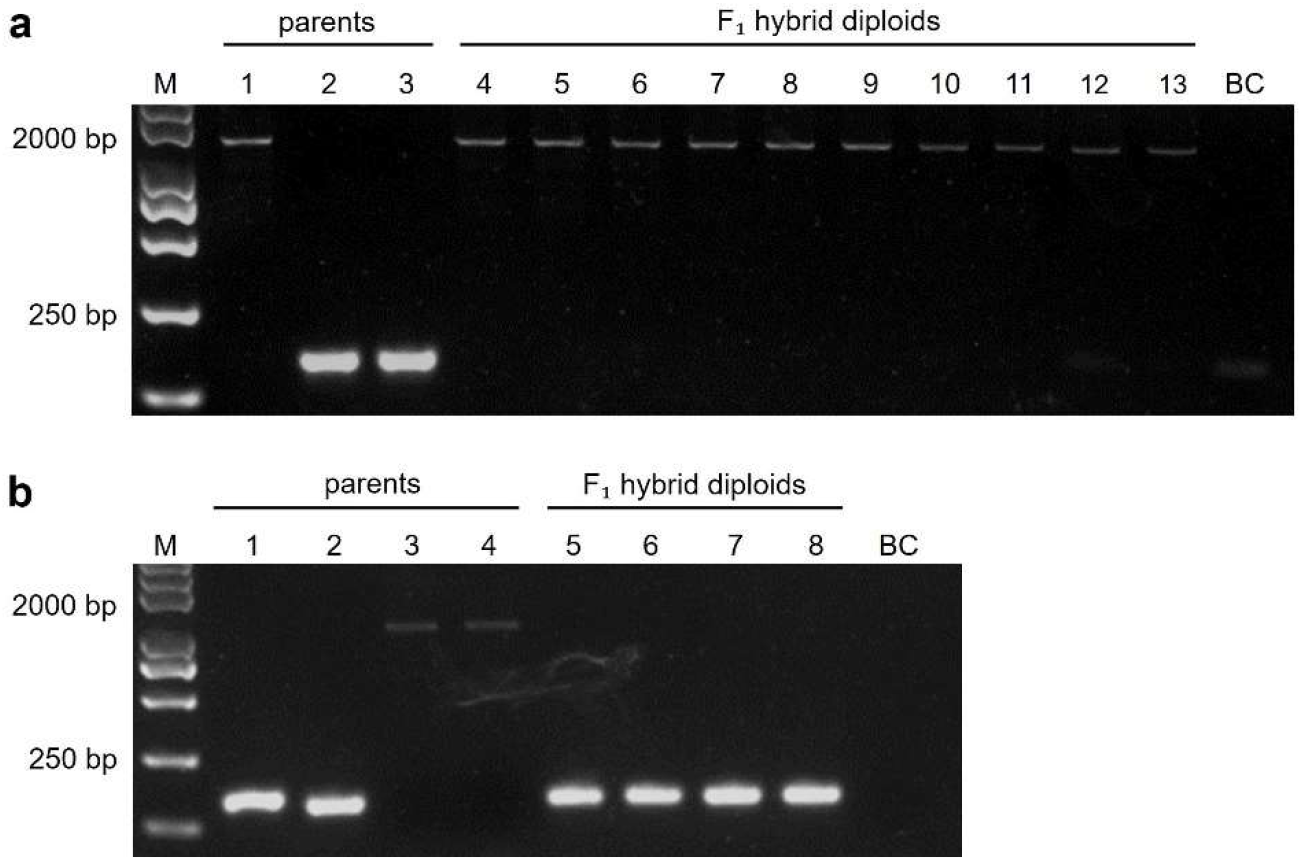
PCR-detection of chloroplast *pet*B marker in P generation and F_1_ hybrid diploids. (a) Parents and F_1_ hybrid diploids in direct cross groups 1 and 2. M: DNA marker; Lane 1: *U. prolifera* mt^−^ parent U687-1; Lane 2–3: *U. linza* mt^+^ parents 23QD07-1 and 23QD07-2; Lane 4–13: F_1_ hybrid diploids. BC: blank control. (b) Parents and F_1_ hybrid diploids in reciprocal cross groups 5 and 6. M: DNA marker; Lane 1–2: *U. linza* mt^−^ parents 24YT12-1 and 23QD07-3; Lane 3–4: *U. prolifera* mt^+^ parent S096-1; Lane 5–8: F_1_ hybrid diploids. BC: blank control.

Second, for the identified F_1_ hybrid diploids, five individuals were selected from each direct cross group and two from each reciprocal cross group; amplification and sequencing results showed that their genotypes and nucleotide sequences both agreed with the paternal parent—*i*.*e*., consistent with the *U. prolifera* mt^−^ parent in direct crosses and with the *U. linza* mt^−^ parent in reciprocal crosses (Fig. 5a, b). Additionally, to determine whether chloroplast mosaicism existed in different parts of the F_1_ hybrid sporophyte thalli, one F_1_ hybrid sporophyte individual was selected from cross group 1 and sampled at the base, middle, tip, and branches of the thallus for detection (Fig. 6a). The results showed that all amplification products matched the *U. prolifera* paternal parent, with no chloroplast mosaicism detected (Fig. 6b). These results demonstrated that, from the P generation to the F_1_ hybrid sporophytes, chloroplasts were inherited in a strictly paternal pattern across both direct and reciprocal crosses, rather than maternal or biparental inheritance.

**Fig. 6.**
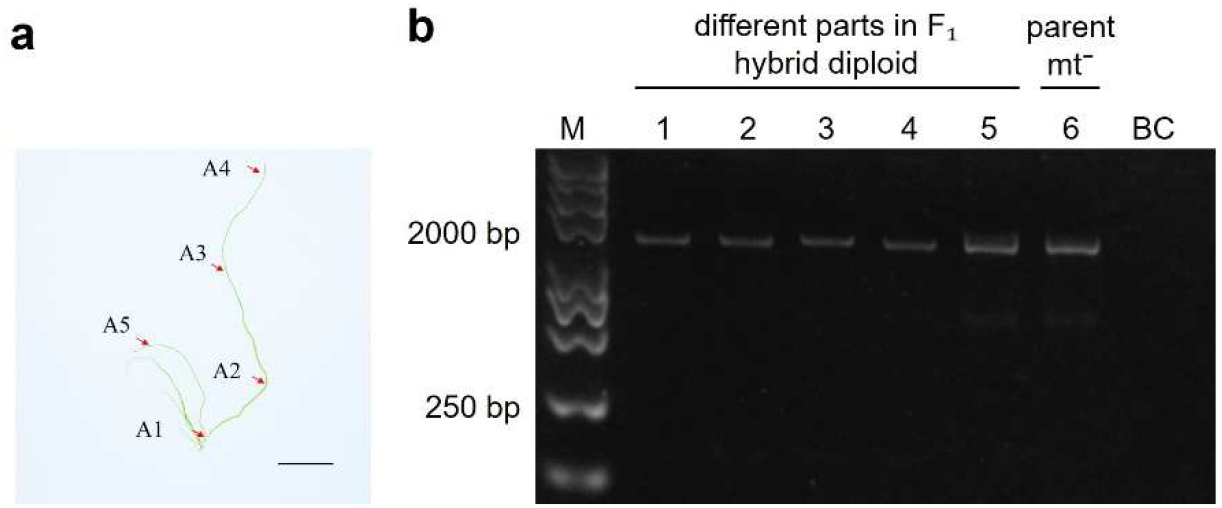
PCR-detection of chloroplast *pet*B marker in different parts of thallus in F_1_ hybrid diploid. (a) M: DNA marker; Lane 1–5: A1– A5; Lane6: *U. prolifera* mt^−^ parent U687-1; BC: blank control. (b) The red arrows point to the sampling parts of a F_1_ hybrid diploid. Scale bar = 1 cm.

Finally, the progeny of F_1_ hybrid sporophytes were also examined. In direct cross group 1, five offspring derived from a single F_1_ hybrid sporophyte—three females and two males—were selected; all yielded *pet*B amplification bands of ∼1.8 kb, matching those of the F_1_ hybrid sporophyte and indicating a paternal origin from *U. prolifera* in the P generation (Fig. 7a). Likewise, in reciprocal cross group 5, five offspring from a single F_1_ hybrid sporophyte—four males and one diploid— were analyzed; all produced *pet*B bands of ∼150 bp, again matching the F_1_ hybrid sporophyte and tracing back to the paternal *U. linza* parent in the P generation (Fig. 7b).

**Fig. 7.**
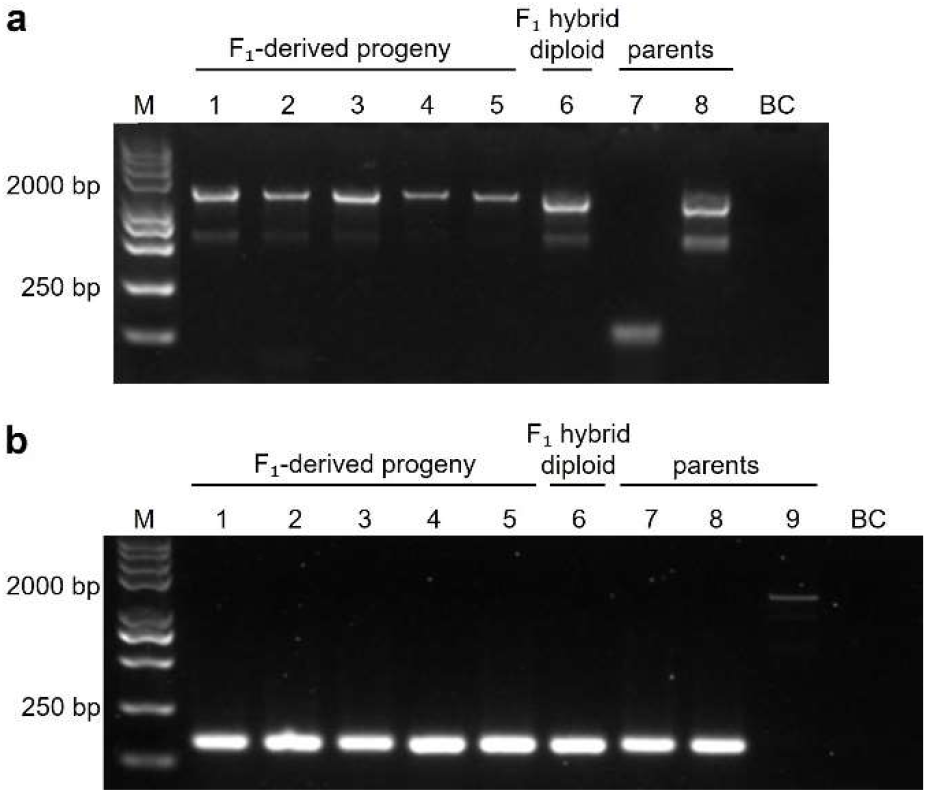
PCR-detection of chloroplast *pet*B marker in F_1_-derived progeny. (a) Parents, F_1_ hybrid diploid and F_1_-derived progeny from direct cross group 1. M: DNA marker; Lane 1–5: F_1_-derived progeny (Lane 2, 4, 5 were mt^+^, and Lane 1, 3 were mt^−^); Lane 6: F_1_ hybrid diploid; Lane 7: *U. linza* mt^+^ parent 23QD07-1; Lane 8: *U. prolifera* mt^−^ parent U687-1; BC: blank control. (b) Parents, F_1_ hybrid diploid and F_1_-derived progeny from reciprocal cross group 5. M: DNA marker; Lane 1–5: F_1_-derived progeny (Lane 1, 3, 4, 5 were mt^−^, and Lane 2 was mt^±^); Lane 6: F_1_ hybrid diploid; Lane 7–8: *U. linza* mt^−^ parent 24YT12-1; Lane 9: *U. prolifera* mt^+^ parent S096-1; BC: blank control.

## 4. Discussion

### 4.1 Bidirectional hybridization compatibility between U. prolifera and U. linza

Our results proved that bidirectional crosses between *U. prolifera* and *U. linza* could both be successful, and the normally developing F_1_ hybrid diploids can be obtained from multiple parental combinations. It should be noted, however, that gametes of *Ulva* possess a strong capacity for parthenogenetic development, and unfertilized gametes may germinate independently to produce haploid progeny (Liu et al., 2015). Therefore, microscopic observation of gamete fusion followed by the development of juvenile thalli is not, by itself, sufficient to exclude interference from parthenogenesis. This represents a common methodological challenge in studies of the *Ulva* life cycle, because *U. prolifera* can produce new individuals through parthenogenesis and other asexual reproductive pathways, making the origin of newly formed thalli difficult to determine (Lin et al., 2008; Gao et al., 2010; Zhang et al., 2016; Liu et al., 2022). The highly differentiated mating-type locus between mt^+^ and mt^−^ strains in *Ulva* was first identified through comparative genomic analyses of *U. partita* (Yamazaki et al., 2017). Subsequently, mating-type-linked molecular markers were developed and validated in *U. prolifera* on the basis of allelic sequence differences within this region (Liu et al., 2022; Xu, 2024). The two markers were detected separately in haploid individuals of the two mating types, occurred together in diploid sporophytes, and segregated in meiotic progeny. This molecular identification system therefore provides a relatively accurate and efficient means of distinguishing true diploid hybrids formed by gamete fusion from haploid individuals arising through parthenogenetic development, thereby improving the reliability of hybrid identification.

Previous studies showed that when artificial hybridization was conducted between *U. prolifera* mt^+^ and *U. linza* mt^−^, gametes rapidly aggregated and fused, whereas in the reciprocal cross, *U. linza* mt^+^ × *U. prolifera* mt^−^, no gamete aggregation was observed, and this combination was considered to exhibit complete prezygotic gametic incompatibility. This asymmetry was previously interpreted as evidence of a certain degree of reproductive separation between the two taxa (Hiraoka et al., 2011, Cui et al., 2018). In contrast, the present study confirmed the successful production of F_1_ hybrid sporophytes (2n) in both the direct and reciprocal interspecific hybridization, indicating the bidirectional hybridization compatibility between *U. prolifera* and *U. linza*. Interestingly, we also observed marked differences in the compatibility between the two hybridization directions; however, the pattern was completely opposite to that reported in previous studies. Specifically, in certain combinations of the direct cross, namely *U. linza* mt^+^ × *U. prolifera* mt^−^, the detection rate of F_1_ diploid hybrid progeny reached 100%; whereas in the reciprocal cross, *U. linza* mt^−^ × *U. prolifera* mt^+^, the maximum detection rate was merely 2.2%–6.6%, with the remaining progeny consisting of haploids formed through parthenogenetic development of parental gametes. Collectively, the results from multiple studies demonstrate that bidirectional interspecific hybridization can occur between *U. prolifera* and *U. linza*, with no absolute reproductive isolation in either direction. The directional bias in hybridization observed in individual experiments is most likely dependent on specific parental combinations, and more precisely, is jointly influenced by factors such as the genetic background and geographic origin of different strains.

### 4.2 Fertility of F_1_ hybrid sporophytes and formation of F_1_-derived haploids

Whether the hybrid progeny can successfully undergo meiosis and produce offspring with normal developmental and reproductive capacity serves as an important criterion for evaluating the fertility of hybrid progeny and its potential genetic and evolutionary significance (Coughlan & Matute, 2020). In this study, molecularly confirmed F_1_ hybrid sporophytes were induced to release reproductive cells, and the mating types for the geminated progeny were subsequently examined. The results showed that the F_1_ hybrid sporophytes not only grew and developed normally, but also formed and released reproductive cells some of which further developed into haploid gametophytes. This demonstrates that the interspecific hybrid progeny possessed the capacity for meiosis and were able to complete the life-cycle transition from F_1_ diploid to haploid. In contrast, some studies have found that, although zygotes could occasionally form in certain combinations between the provisionally named “*U. flexuosa*” clade and *U. californica*, the progeny mostly arrested at early developmental stages or failed to complete meiosis to release normal spores, indicating that the formation of hybrid zygotes does not necessarily imply complete hybrid compatibility (Hiraoka et al., 2017). Notably, F_1_ hybrid diploids derived from our reciprocal cross, upon maturation, could either produce diploid spores for asexual reproduction or generate gametophytes—though exclusively male—suggesting that hybridization may have severely interfered with normal meiotic progression. This provided a typical example of hybrid apomicts in algae (Kamiya et al., 2011). Through continuous molecular assessment of mating types, the present study established a relatively complete line of evidence spanning gamete fusion, diploid formation, meiosis, and the production of haploid progeny. These findings indicate that *U. prolifera* and *U. linza* are capable of producing fertile recombinant offspring in specific parental combinations. Because meiosis reshuffles genetic material from both parents, entry of the hybrid progeny into the haploid phase may generate a range of genetically distinct strains, thereby providing a basis for subsequent natural selection, backcrossing with parental taxa, and potential introgression (Twyford & Ennos, 2012; Suarez-Gonzalez et al., 2018). The recovery of hybrid progeny capable of completing meiosis in specific crosses therefore provides experimental support for further assessment of hybrid compatibility between *U. prolifera* and *U. linza* and for subsequent genetic studies.

### 4.3 The paternal inheritance of chloroplasts in hybrid offspring

In the present study, we demonstrated that chloroplasts in *U. prolifera* × *U. linza* hybrids exhibited a stable paternal inheritance pattern. This inheritance mode was consistently observed in both direct and reciprocal crosses, in different thallus regions of F_1_ hybrid sporophytes, and in F_1_-derived haploid progeny. No evidence of biparental chloroplast retention, chloroplast chimerism, or subsequent segregation of chloroplast types was detected. These results suggest that chloroplast inheritance in this interspecific hybrid system is likely determined at an early stage after fertilization and can be stably maintained throughout vegetative development and life-cycle transition. Our findings provide new insights into the flexibility of organelle inheritance in *Ulva* and highlight the potential influence of interspecific hybridization on cytoplasmic inheritance patterns.

During fertilization, chloroplasts contributed by both parental gametes may coexist transiently in the zygote (Birky, 1995). However, selective mechanisms, including differential degradation of parental chloroplasts, differences in organelle replication efficiency, and biased partitioning during cell division, may determine the relative contribution of parental chloroplast genomes and ultimately result in maternal, paternal, or biparental inheritance patterns (Miyamura, 2010; Sakamoto & Takami, 2024). Maternal inheritance represents the predominant chloroplast transmission mode in most land plants and green algae, although paternal and biparental inheritance patterns have also been reported in several lineages (Mogensen, 1996; Miyamura, 2010). Within the genus *Ulva*, early studies based on autoradiographic observations suggested paternal chloroplast inheritance in *U. mutabilis* (Bråten, 1973), the synonym of *U. compressa* (Steinhagen et al., 2019). However, recent molecular marker-based analyses indicated predominantly maternal chloroplast inheritance in *U. compressa* (Kagami et al., 2008). Further investigations involving a large number of intraspecific hybrid progenies revealed that, although maternal inheritance was overwhelmingly dominant, rare cases of paternal inheritance or biparental chloroplast retention could also occur (Mogi et al, 2009). For the two parental species examined in the present study, previous analyses based on crosses between different populations of *U. prolifera* demonstrated maternal chloroplast inheritance (Liu, 2022), whereas chloroplast inheritance patterns in *U. linza* remain unexplored. Given the close genetic relationship between *U. prolifera* and *U. linza*, similar maternal inheritance patterns would generally be expected in these species.

Our study provides the first evidence of chloroplast inheritance patterns in interspecific *Ulva* hybrids, revealing stable paternal chloroplast transmission in *U. prolifera* × *U. linza* progeny. From this, we make a bold hypothesis that in the offspring resulting from *U. prolifera* × *U. linza*, the chloroplast inheritance patterns has reversed compared to that of the parent species. Similar shifts in chloroplast inheritance patterns have been documented in several terrestrial plant groups, including *Cicer, Actinidia*, and *Passiflora*, where hybrids exhibited paternal or biparental chloroplast inheritance despite predominantly maternal inheritance in parental species (Kumari et al., 2011; Li et al., 2013; Shrestha et al., 2021). These examples indicated that organelle inheritance can be modulated by specific hybrid combinations and genetic backgrounds rather than being strictly conserved at the species level (Burton RS, 2013). To our knowledge, such a hybrid-associated alteration of chloroplast inheritance has not previously been reported in algae. Therefore, we propose that interspecific hybridization between *U. prolifera* and *U. linza* may modify the selective processes governing chloroplast transmission, resulting in preferential retention of paternal chloroplasts. Several mechanisms may potentially explain this unusual inheritance pattern. These include a greater contribution of chloroplasts from mt^−^ gametes during fertilization, selective elimination of mt^+^-derived chloroplasts after zygote formation, or differential competitive abilities of parental chloroplast genomes during replication and intracellular segregation (Birky, 1995; Kagami et al., 2008; Miyamura, 2010). However, the present study mainly inferred chloroplast origin based on molecular markers, and direct cytological evidence during early fertilization stages is still lacking. Future studies should first determine the chloroplast inheritance pattern of *U. linza* through controlled intraspecific crosses. In addition, fluorescence-based chloroplast tracing, ultrastructural observations, and time-course analyses following fertilization will be necessary to clarify the dynamic processes underlying paternal chloroplast retention and maternal chloroplast exclusion in hybrids of *U. prolifera* × *U. linza*.

### 4.4 Ecological and evolutionary implications

The present study revealed that the *U. prolifera* × *U. linza* F_1_ hybrid sporophytes inherited paternal chloroplasts, and could undergo normal meiosis upon maturation, which bears significant ecological and evolutionary implications. First, the completion of meiosis indicates that genetic materials from the two species can potentially be transmitted to subsequent generations through continuous reproduction or backcrossing. Consequently, gene flow between traditionally recognized species may occur in regions where they coexist, leading to genetic introgression and alterations in the genetic composition of local populations. Second, chloroplasts play essential roles in photosynthesis, energy conversion, and various metabolic processes (Hoefnagel et al., 1998; Song et al., 2021). The combination of a paternal chloroplast genome with a recombinant nuclear background may generate novel cytonuclear interactions, potentially influencing offspring growth performance, environmental tolerance, and ecological fitness (Burton et al., 2013). Stable paternal chloroplast inheritance further suggested that the chloroplast lineage of the mt^−^ parent could be preferentially retained and propagated in hybrid populations. This biased transmission may result in discordance between chloroplast-based phylogenetic relationships and nuclear genomic relationships, which should be carefully considered when chloroplast markers were used for species identification, population tracing, or phylogeographic analyses.

In the Yellow Sea area, *U. prolifera* and *U. linza* shared extensive overlapping habitats including the *Pyropia* aquaculture rafts (Han et al., 2020). They both appeared simultaneously at the early stage of the Yellow Sea green tide (Zhang et al., 2019), and released a large number of microspores during the course of the green tide. This created abundant opportunities for interspecific hybridization between the two species in their natural environment. Such processes may increase genetic diversity within *Ulva* populations and generate novel genotypes with altered adaptive capacity, reproductive performance, or dispersal potential (Fort et al., 2021), thereby potentially influencing community succession and the formation of dominant bloom-forming populations.

Therefore, ecological monitoring and risk assessment of green tides should consider not only species composition and biomass dynamics but also reproductive compatibility among closely related taxa, the potential frequency of natural hybridization, and the direction of organelle lineage transmission. Meanwhile, the hybrid system provides a valuable experimental model for investigating species diversification, cytonuclear coevolution, and the genetic improvement of economically and ecologically important algal resources.

## Abbreviations

5S-NTS: 5S rDNA non-transcribed spacer
ITS: internal transcribed spacer
VSE medium: von Stosch’s enriched medium

## Credit Author Statement

**Zhengzhao Xu:** Methodology, Validation, Investigation, Formal analysis, Visualization, Data curation; Writing - original draft. **Jin Zhao**: Methodology, Resources. **Peng Jiang:** Conceptualization, Funding acquisition, Supervision, Project administration, Writing - review & editing.

## Acknowledgements

This research was funded by the Science & Technology Basic Resources Investigation Program of China (2018FY100205), Shandong Provincial Natural Science Foundation (ZR2024MD028), Natural Science Foundation of Qingdao City (23-2-1-172-zyyd-jch), and Key R&D Program of Shandong Province (2019GSF107012).

## Statement regarding informed consent, human/animal rights

No conflicts, informed consent, or human or animal rights are applicable to this study.

## References

Birky CW. Uniparental inheritance of mitochondrial and chloroplast genes: mechanisms and evolution. Proceedings of the National Academy of Sciences of the United States of America, 1995, 92: 11331–11338. 10.1073/pnas.92.25.11331.

Bråten T. Autoradiographic evidence for the rapid disintegration of one chloroplast in the zygote of the green alga Ulva mutabilis. Journal of Cell Science, 1973, 12: 385–389. 10.1242/jcs.12.2.385.

Burton RS, Pereira RJ, Barreto FS. Cytonuclear genomic interactions and hybrid breakdown. Annual Review of Ecology, Evolution, and Systematics, 2013, 44: 281–302. 10.1146/annurev-ecolsys-110512-135758.

Choi JW, Graf L, Peters AF, Cock JM, Nishitsuji K, Arimoto A, Shoguchi E, Nagasato C, Choi CG, Yoon HS. Organelle inheritance and genome architecture variation in isogamous brown algae. Scientific Reports, 2020, 10: 2048. 10.1038/s41598-020-58817-7.

Coughlan JM, Matute DR. The importance of intrinsic postzygotic barriers throughout the speciation process. Philosophical Transactions of the Royal Society B: Biological Sciences, 2020, 375: 20190533. 10.1098/rstb.2019.0533.

Cui JJ, Monotilla AP, Zhu WR, Takano Y, Shimada S, Ichihara K, Matsui T, He PM, Hiraoka M. Taxonomic reassessment of Ulva prolifera (Ulvophyceae, Chlorophyta) based on specimens from the type locality and Yellow Sea green tides. Phycologia, 2018, 57: 692–704. 10.2216/17-139.1.

Duong TT, Nguyen HTT, Nguyen HT, Nguyen QT, Nguyen BD, Chuong NN, Chu HD, Tran LSP. Advances in the genus Ulva research: From structural diversity to applied utility. Plants, 2025, 14: 3052. 10.3390/plants14193052.

Fort A, McHale M, Cascella K, Potin P, Usadel B, Guiry MD, Sulpice R. Foliose Ulva species show considerable inter-specific genetic diversity, low intra-specific genetic variation, and the rare occurrence of inter-specific hybrids in the wild. Journal of Phycology, 2021, 57: 219–233. 10.1111/jpy.13079.

Gao S, Chen XY, Yi QQ, Wang GC, Pan GH, Lin AP, Peng G. A strategy for the proliferation of Ulva prolifera, main causative species of green tides, with formation of sporangia by fragmentation. PLoS ONE, 2010, 5: e8571. 10.1371/journal.pone.0008571.

Han HB, Li Y, Fan Sl, Song W, Li Y, Xiao J, Wang ZL, Zhang XL, Ding DW. The contribution of attached Ulva prolifera on Pyropia aquaculture rafts to green tides in the Yellow Sea. Acta Oceanologica Sinica, 2020, 39: 101–106. 10.1007/s13131-019-1452-0.

Hiraoka M, Dan A, Shimada S, Hagihira M, Migita M, Ohno M. Different life histories of Enteromorpha prolifera (Ulvales, Chlorophyta) from four rivers on Shikoku Island, Japan. Phycologia, 2003, 42: 275–284. 10.2216/i0031-8884-42-3-275.1.

Hiraoka M, Ichihara K, Zhu WR, Ma JH, Shimada S. Culture and hybridization experiments on an Ulva clade including the Qingdao strain blooming in the Yellow Sea. PLoS ONE, 2011, 6(5): e19371. 10.1371/journal.pone.0019371.

Hiraoka M, Ichihara K, Zhu WR, Shimada S, Oka N, Cui JJ, Tsubaki S, He PM. Examination of species delimitation of ambiguous DNA-based Ulva (Ulvophyceae, Chlorophyta) clades by culturing and hybridisation. Phycologia, 2017, 56: 517–532. 10.2216/16-109.1.

Hoefnagel MHN, Atkin OK, Wiskich JT. Interdependence between chloroplasts and mitochondria in the light and the dark. Biochimica et Biophysica Acta (BBA)-Bioenergetics, 1998, 1366: 235–255. 10.1016/S0005-2728(98)00126-1.

Hofmann LC, Nettleton JC, Neefus CD, Mathieson AC. Cryptic diversity of Ulva (Ulvales, Chlorophyta) in the Great Bay Estuarine System (Atlantic USA): introduced and indigenous distromatic species. European Journal of Phycology, 2010, 45: 230–239. 10.1080/09670261003746201.

Ichihara, K., Yamazaki T, Miyamura S, Hiraoka M, Kawano S. Asexual thalli originated from sporophytic thalli via apomeiosis in the green seaweed Ulva. Scientific Reports, 2019, 9: 1–12. 10.1038/s41598-019-50070-x.

Innes DJ. Genetic structure of asexually reproducing Enteromorpha linza (Ulvales: Chlorophyta) in Long Island Sound. Marine Biology, 1987, 94: 459–467. 10.1007/BF00428253.

Kagami Y, Mogi Y, Arai T, Yamamoto M, Kuwano K, Kawano S. Sexuality and uniparental inheritance of chloroplast DNA in the isogamous green alga Ulva compressa (Ulvophyceae). Journal of Phycology, 2008, 44: 691–702. 10.1111/j.1529-8817.2008.00527.x.

Kamiya M, West JA, Hara Y. Induction of apomixis by outcrossing between genetically divergent entities of Caloglossa leprieurii (Ceramiales, Rhodophyta) and evidence of hybrid apomicts in nature. Journal of Phycology, 2011, 47: 753–762. 10.1111/j.1529-8817.2011.01016.x.

Kang JH, Jang JE, Kim JH, Byeon SY, Kim S, Choi SK, Kang YH, Park SR, Lee HJ. Species composition, diversity, and distribution of the genus Ulva along the coast of Jeju Island, Korea based on molecular phylogenetic analysis. PLoS ONE, 2019, 14: e0219958. 10.1371/journal.pone.0219958.

Kumari M, Clarke HJ, Colas des Francs-Small C, Small I, Khan TN, Siddique KHM. Albinism does not correlate with biparental inheritance of plastid DNA in interspecific hybrids in Cicer species. Plant Science, 2011, 180: 310–316. 10.1016/j.plantsci.2011.01.003.

Leskinen E, Pamilo P. Evolution of the ITS sequences of ribosomal DNA in Enteromorpha (Chlorophyceae). Hereditas, 1997, 126: 17–23. 10.1111/j.1601-5223.1997.00017.x.

Li D, Qi X, Li X, Li L, Zhong C, Huang H. Maternal inheritance of mitochondrial genomes and complex inheritance of chloroplast genomes in Actinidia Lindl.: evidences from interspecific crosses. Molecular Genetics and Genomics, 2013, 288: 101–110. 10.1007/s00438-012-0732-6.

Lin AP, Shen SD, Wang JW, Yan BL. Reproduction diversity of Enteromorpha prolifera. Journal of Integrative Plant Biology, 2008, 50: 622–629. 10.1111/j.1744-7909.2008.00647.

Liu Q, Yu RC, Yan T, Zhang QC, Zhou MJ. Laboratory study on the life history of bloom-forming Ulva prolifera in the Yellow Sea. Estuarine, Coastal and Shelf Science, 2015, 163: 82–88. 10.1016/j.ecss.2014.08.011.

Liu QC, Wu CH, Xie WF, Zhao J, Jiang P. Validation of mating type-related markers in Ulva prolifera (Ulvophyceae, Chlorophyta) and their detection during various reproductive modes, Algal Research, 2022a, 62: 102611. 10.1016/j.algal.2021.102611.

Liu WZ. Comparative genomic analysis on the chloroplast of Ulva prolifera and developments of molecular markers. Master thesis, University of Chinese Academy of Sciences, 2022. (in Chinese with English abstract)

Liu WZ, Liu QC, Zhao J, Wei X, Jiang P. Comparative chloroplast genomes of Ulva prolifera and U. linza (Ulvophyceae) provide genetic resources for the development of interspecific markers. Journal of Oceanology and Limnology, 2022b, 40: 2372–2384. DOI: 10.1007/s00343-022-2045-x.

Miyamura, S. Cytoplasmic inheritance in green algae: patterns, mechanisms and relation to sex type. Journal of Plant Research, 2010, 123: 171–184. 10.1007/s10265-010-0309-6.

Mogensen HL. The hows and whys of cytoplasmic inheritance in seed plants. American Journal of Botany, 1996, 83: 383–404. 10.2307/2446175.

Mogi Y, Hatakeyama Y, Kuwano K, Miyamura S, Kawano S. Patterns of organelle inheritance revealed by 12 interline crosses in Ulva compressa. Phycologia, 2009, 48(Suppl): 86.

Reboud X, Zeyl C. Organelle inheritance in plants. Heredity, 1994, 72: 132–140. 10.1038/hdy.1994.19.

Sakamoto W, Takami T. Plastid inheritance revisited: emerging role of organelle DNA degradation in angiosperms. Plant and Cell Physiology, 2024, 65: 484–492. 10.1093/pcp/pcad104.

Saunders GW, Kucera H. An evaluation of rbcL, tufA, UPA, LSU and ITS as DNA barcode markers for the marine green macroalgae. Cryptogamie, Algologie, 2010, 31: 487–528.

Shimada S, Yokoyama N, Arai S, Hiraoka M. Phylogeography of the genus Ulva (Ulvophyceae, Chlorophyta), with special reference to the Japanese freshwater and brackish taxa. Journal of Applied Phycology, 2008, 20: 979–989. 10.1007/s10811-007-9296-y.

Shrestha B, Gilbert LE, Ruhlman TA, Jansen RK. Clade-specific plastid inheritance patterns including frequent biparental inheritance in Passiflora interspecific crosses. International Journal of Molecular Sciences, 2021, 22: 2278. 10.3390/ijms22052278.

Sloan DB, Warren JM, Williams AM, Wu Z, Abdel-Ghany SE, Chicco AJ, Havird JC. Cytonuclear integration and coevolution. Nature Reviews Genetics, 2018, 19: 635–648. 10.1038/s41576-018-0035-9.

Song Y, Feng L, Alyafei MAM, Jaleel A, Ren M. Function of chloroplasts in plant stress responses. International Journal of Molecular Sciences, 2021, 22: 13464. 10.3390/ijms222413464.

Steinhagen S, Barco A, Wichard T, Weinberger F. Conspecificity of the model organism Ulva mutabilis and Ulva compressa (Ulvophyceae, Chlorophyta). Journal of Phycology, 2019, 55: 25–36. 10.1111/jpy.12804.

Steinhagen S, Karez R, Weinberger F. Cryptic, alien and lost species: molecular diversity of Ulva sensu lato along the German coasts of the North and Baltic Seas. European Journal of Phycology, 2019, 54: 466–483. 10.1080/09670262.2019.1597925.

Suarez-Gonzalez A, Lexer C, Cronk QCB. Adaptive introgression: a plant perspective. Biology Letters, 2018, 14: 20170688. 10.1098/rsbl.2017.0688.

Tran LAT, Vieira C, Steinhagen S, Maggs CA, Hiraoka M, Shimada S, Nguyen TV, De Clerck O, Leliaert F. An appraisal of Ulva (Ulvophyceae, Chlorophyta) taxonomy. Journal of Applied Phycology, 2022, 34: 2689–2703. 10.1007/s10811-022-02815-x.

Twyford AD, Ennos RA. Next-generation hybridization and introgression. Heredity, 2012, 108: 179–189. 10.1038/hdy.2011.68.

Xie EY, Xu RS, Zhang JH, Xu C, Huang BW, Zhu WR, Cui JJ. Growth characteristics of hybrids produced by closely related Ulva species. Aquaculture, 2020, 519: 734902. 10.1016/j.aquaculture.2019.73490.

Xie WF, Wu CH, Zhao J, Lin XY, Jiang P. New records of Ulva spp. (Ulvophyceae, Chlorophyta) in China, with special reference to an unusual morphology of U. meridionalis forming green tides. European Journal of Phycology, 55: 412–425. 10.1080/09670262.2020.1740946.

Xu H. Expression characteristics of ACTIN gene in two species of seaweeds and development of its regulatory elements and molecular markers. Master thesis, University of Chinese Academy of Sciences, 2024. (in Chinese with English abstract)

Yamazaki T, Ichihara K, Suzuki R, Oshima K, Miyamura S, Kuwano K, Toyoda A, Suzuki Y, Sugano S, Hattori M, Kawano S. Genomic structure and evolution of the mating type locus in the green seaweed Ulva partita. Scientific Reports, 2017, 7: 11679. 10.1038/s41598-017-11677-0.

Zhang JH, Kim JK, Yarish C, He PM. The expansion of Ulva prolifera O.F. Müller macroalgal blooms in the Yellow Sea, PR China, through asexual reproduction. Marine Pollution Bulletin, 2016, 104: 101–106. 10.1016/j.marpolbul.2016.01.056.

Zhang JH, Shi JT, Gao S, Huo YZ, Cui JJ, Shen H, Liu GY, He PM. Annual patterns of macroalgal blooms in the Yellow Sea during 2007–2017. PLoS ONE, 2019, 14: e0210460. 10.1371/journal.pone.0210460.

Zhang Q Sodmergen. Why does biparental plastid inheritance revive in angiosperms? Journal of Plant Research, 2010, 123: 201–206. 10.1007/s10265-009-0291-z.

